# Dimensionality Reduction and Denoising of Spatial Transcriptomics Data Using Dual-Channel Masked Graph Autoencoder

**DOI:** 10.1101/2024.05.30.596562

**Authors:** Wenwen Min, Donghai Fang, Jinyu Chen, Shihua Zhang

## Abstract

Recent advances in spatial transcriptomics (ST) technology allow researchers to comprehensively measure gene expression patterns at the level of individual cells or even subcellular compartments while preserving the spatial context of their tissue. Spatial domain identification is a critical task in analyzing the ST data. However, effectively capturing distinctive gene expression features and relationships between genes poses a significant challenge. We develop a graph self-supervised learning method STMask for the analysis and exploration of the ST data. STMask combines the masking mechanism with a graph autoencoder, compelling the gene representation learning channel to acquire more expressive representations. Simultaneously, it combines the masking mechanism with graph self-supervised contrastive learning methods, pulling together the embedding distances between spatially adjacent points and pushing apart the representations of different clusters, allowing the gene relationship learning channel to learn more comprehensive relationships. The applications of STMask to four ST datasets demonstrate that STMask outperforms state-of-the-art methods in various tasks, including spatial clustering and trajectory inference. Source code is available at https://github.com/donghaifang/STMask.

**Author summary:** Spatial Transcriptomics (ST) is an emerging transcriptomic sequencing technology aimed at revealing the spatial distribution of gene expression and cell types within tissues. This method enables the acquisition of gene expression profiles at the level of individual cells or spots within the tissue, uncovering the spatial expression patterns of genes. However, accurately identifying spatial domains in ST data remains challenging. In our study, we introduce STMask, a self-supervised learning method that combines a dual-channel masked graph autoencoder with masking and contrastive learning. Our work contributes primarily in two aspects: (1) We propose a novel graph self-supervised learning method (STMask) specifically tailored for the analysis and research of ST data, which enhances the ability to capture the unique features of gene expression and spatial relationships within tissues. (2) Through comprehensive experiments, STMask provides valuable insights into biological processes, particularly in the context of breast cancer. It identifies enrichment of various differentially expressed genes in tumor regions, such as *IGHG1*, which can serve as effective targets for cancer therapy.

## Introduction

Within complex organisms, cells are organized into similar clusters within spatial context [1, 2]. The intricate arrangement of tissues mirrors the dynamic interactions and specialized functions among cells, where the precise positional cues of transcriptional expressions play a pivotal role in unraveling biological functions and networks [3, 4]. The latest ST technologies, such as 10x Visium [5] and Stereo-seq [6], enable whole-genome analysis at the resolution of cellular or even subcellular levels, capturing gene expression corresponding to specific locations known as spots. These spatial insights provide a solid foundation for understanding many biological processes influencing disease.

Identifying spatial domains (regions with similar spatial expression patterns) is crucial in analyzing ST data to understand expression patterns [7]. Currently, spatial domain identification primarily involves non-spatial and spatial clustering methods [1]. Traditional non-spatial clustering methods, like the Louvain algorithm, only use gene expression data as input. This limitation can lead to disjointed clustering results within tissue sections. These methods do not effectively utilize spatial information, resulting in clustering outcomes that poorly align with tissue sections.

To address this challenge, some spatial clustering methods have recently been proposed [3, 8]. These methods consider the similarity between neighboring spots by leveraging spatial positional information, aiming to elucidate the spatial dependence of gene expression. stLearn [9] employs Louvain clustering to process graph adjacency matrices, effectively integrating gene expression with spatial information and histological morphological features for spatial clustering. Additionally, BayesSpace [10] utilizes a fully Bayesian statistical model to optimize spatial neighboring spot relationships to reflect the spatial structure and identify cells. Despite the potential of spatial positional information to enhance clustering accuracy, the aforementioned spatial clustering algorithms have yet to achieve optimal performance [3].

Graph Neural Networks (GNNs) have shown their effectiveness in integrating gene expression and spatial positioning [11, 12]. These models learn low-dimensional embeddings within the GNN framework, capturing expressive features while preserving spot characteristics and topological structure. To leverage the advantages of GNNs in ST data, several GNN-based models have been proposed for spatial domain identification [13]. For instance, SpaGCN [14] combines Graph Convolutional Networks (GCNs) with gene expression, histological image, and spatial positional information to identify variable genes in spatial domains. It utilizes unsupervised clustering algorithms to detect different spatial expression patterns.

Moreover, some approaches for analyzing ST data are based on graph auto-encoding or graph contrastive learning to acquire latent representations [15]. These representations are subsequently employed in downstream task analysis. For instance, the spatial clustering method DeepST [16], based on graph auto-encoding, takes gene expression, histological images, and spatial information as input. It employs GCN as encoders and reconstructs the input graph topology and spot features to detect and identify spatial domains. SEDR [17] adopts a deep auto-encoder network to learn gene representations and utilizes a variational graph auto-encoder to embed gene and spatial information simultaneously. CCST [18] designed based on graph contrastive learning integrates spatial structure and gene expression information by stacking multiple layers of GCN. It utilizes the concept of DGI [19] to construct positive and negative pairs to facilitate ST data analysis. STAGATE [1] integrates spatial information and gene expression using adaptive graph attention auto-encoders to enhance the accuracy of spatial domain identification. However, these methods either overly rely on global graph information, resulting in the misuse of redundant graph structure information during training, thereby limiting performance and failing to attain more robust representations, or they overlook the local correlations between spots, leading to an inability to acquire effective discriminative power.

Considering the above two problems, we propose a graph self-supervised learning method STMask consisting of two channels for the analysis and exploration of ST data with the masking mechanism. The first channel is dedicated to gene representation learning, employing a graph auto-encoding approach with masking to compel the model to extract more expressive gene features from neighboring spots that have not been masked. The second channel, termed gene relationship learning, incorporates two distinct topological views under masking, effectively capturing discriminative relationships between spots within a contrastive learning framework. To assess the effectiveness of STMask, we tested its spatial clustering performance on four different datasets. The experiments demonstrated that STMask achieved competitiveness compared to existing methods. STMask could also identify various differentially expressed genes, such as *IGHG1*, which could serve as effective targets for cancer therapy, displaying its great power.

## Materials and methods

### Dataset description

In this study, we analyze four different datasets Table 1. We first analyze the 10x Vision dataset, which consists of 12 sections of the human dorsolateral prefrontal cortex (DLPFC) containing spatial expression information. Maynard et al. [21] have manually annotated the layers of the DLPFC and white matter (WM) based on morphological features and gene markers. Each section has five to seven regions that are manually annotated (Table A in S1 Supplementary materials).

**Table 1.** The statistics of the datasets used in this study.

| Datasets | Spots | Slices | Genes | Domains | Platforms |
| --- | --- | --- | --- | --- | --- |
| DLPFC [21] | 3,460-4,789 | 12 | 33,538 | 5-7 | 10x Visium |
| BRCA [17] | 3,798 | 1 | 36,601 | 20 | 10x Visium |
| HM [22] | 293 | 1 | 16,148 | 4 | Spatial Transcriptomics |
| MBA [5] | 2,695 | 1 | 32,285 | 52 | 10x Visium |

In the second dataset, we analyze the human breast cancer (BRCA) dataset from 10x Vision, which comprises 3,798 spots and 36,601 genes. The histological images of this dataset exhibit distinct regional shapes. Xu et al. [17] manually annotate the dataset, covering 20 regions.

The third dataset originates from human melanoma (HM) samples from the Spatial Transcriptomics platform. It comprises 293 points and 16,148 genes. Thrane et al. [22] manually annotate three distinct regions: melanoma, stroma, and lymphoid tissue, along with an additional unannotated region. Hence, we utilize these four domains to assess spatial domain recognition performance.

Finally, we analyze the anterior of the mouse brain tissue (MBA) dataset from 10x Visium, comprising 2,695 spots and 32,285 genes [5]. The Allen Mouse Brain Reference Atlas annotates this dataset into 51 known clusters and one unmarked region manually [20]. Thus, we consider 52 clusters as recognizable spatial domains.

### Data preprocessing and spatial graph construction

STMask filters ST data by retaining genes present in the transcriptome at a minimum of 50 locations with a count of at least 10 at each spot. After normalization, highly variable genes are selected for denoising tasks. Principal Component Analysis (PCA) reduces dimensionality, with the top *N*_*f*_ principal components forming the initial gene expression matrix 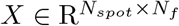, where *N*_*spot*_ is the number of spots.

The key advantage of ST data lies in its use of spatial information to probe relationships between neighboring spots. Leveraging the “homophily” principle, akin individuals tend to connect in social networks or communities. Employing the K-nearest neighbors (KNN) algorithm, we compute Euclidean distances between spots, assuming each spot’s K nearest neighbors share its characteristics [23].

Therefore, we construct an initial undirected and unweighted topological graph 𝒢 = (𝒱, ℰ), where 𝒱 = {*v*_*i*_} represents the set of vertices corresponding to spots, and ℰ is the corresponding set of edges. Specifically, each spot vertex *v*_*i*_ is associated with a *N*_*f*_ -dimensional feature vector 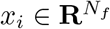, forming the feature vector set *X* = {*x*_*i*_}, where *X* serves as the initialized gene expression matrix. The adjacency matrix 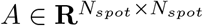 is defined by the edge set ℰ : if spot *j* is a neighboring node of spot *i*, then *A*_*ij*_ = 1; otherwise, *A*_*ij*_ = 0.

### Data augmentation

When input features match or are fewer than encoding dimensions, unconstrained vanilla graph autoencoders can suffer from the notorious “identity function” problem [24], merely copying input to output without learning useful information. Denoising autoencoders intentionally corrupt input, preventing such trivial mapping and encouraging the decoder to extract valuable information from noisy data [25]. Augmenting input with masks and graph structures addresses this issue effectively.

We set a hyperparameter *ρ*_*m*_ for the feature mask rate, then randomly select a subset 𝒱_*m*_ from spots 𝒱 to mask. This creates an augmented attribute graph, denoted 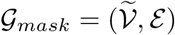, where 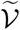 is the vertex set of all spots, but some gene expression matrix values are replaced by the mask token [MASK], represented by 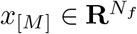. Formally, the augmented feature representation 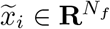 for any spot in 𝒢_*mask*_ is as follows:

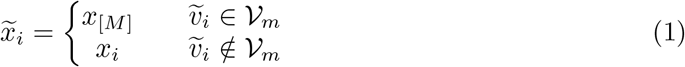

Similarly, a straightforward and effective way to create another masking graph is by sampling a subset ℰ_*drop*_ from the edge set ℰ using a predetermined edge mask rate *ρ*_*d*_, e.g., Bernoulli distribution: ℰ_*drop*_ *∼ Bernoulli*(*ρ*_*d*_). Consequently, the input for the gene relationship learning channel is the masked graph 𝒢_*drop*_ = (𝒱, ℰ_*vis*_). Here, we have ℰ_*vis*_ ∪ ℰ_*drop*_ = ℰ and ℰ_*vis*_ ∩ ℰ_*drop*_ = ∅, where ℰ_*vis*_ represents the retained set of edges used for generating latent representations. The sampled adjacency matrix *Ã* is defined by the edge set ℰ_*vis*_.

### Gene representation learning channel based on graph autoencoder model

Graph convolutional networks demonstrate robust capabilities in processing and aggregating information from neighboring nodes. Therefore, as previously mentioned, to facilitate gene representation learning, we utilize GCN as the backbone model and construct an autoencoder comprising an encoder and a decoder. The input consists of the enhanced gene expression matrix 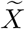 with feature masking and the adjacency matrix *A* capturing spatial location relationships, referred to as 𝒢_*mask*_. Through the encoder, we map this input into a more robust latent representation 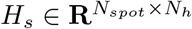, where *N*_*h*_ is the feature dimension of the latent representation, and then the decoder reconstructs the masked spots from the unmasked spots, resulting in the output 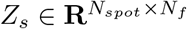.

Specifically, the objective of the encoder (consists of multiple layers of GCN) is to learn a mapping function ℱ^*en*^ under the parameter *θ*, such that it satisfies the following criterion:

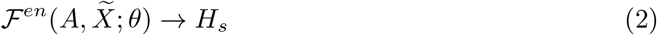

After the model training is completed, we take the initial gene attribute graph 𝒢 as input (without any masking operations) and utilize the output *H* from the encoder for downstream analysis tasks of ST data.

To enhance the robustness of the autoencoder, we employ the re-mask technique to process the learned *H*_*s*_ before feeding them into the decoder. This technique compels the graph decoder to extract valuable information from adjacent spots that have not been masked. In terms of form, we use the masking token [DMASK], i.e., a decoder mask vector 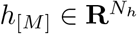, to substitute certain feature vectors of the latent representation *H*_*s*_. Therefore, we take the re-masked 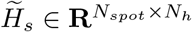 as the input to the graph decoder, which is defined as follows:

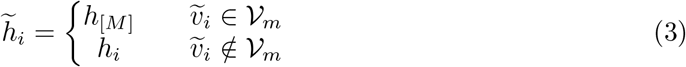

Under the parameter *ϕ*, the decoder, composed of multiple layers of GCN, learns a mapping function ℱ^*de*^ to reconstruct the latent representation into the initial gene expression matrix *X*. The process is as follows:

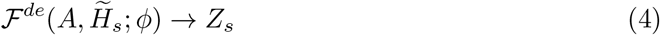

### Gene relationship learning channel based on graph contrastive learning

Based on the assumption of homogeneity, the self-supervised graph contrastive learning method strengthens the connections between adjacent spots in the relational network and separates spots from different clusters. To ensure highly discriminative relationships in the model, adopting a feasible contrastive learning method is a natural choice. This method requires maximizing mutual information from multiple perspectives.

Therefore, in the gene relationship learning channel, we use the masked graph 𝒢_*drop*_ as input and leverage the encoder shared with the gene representation learning channel to learn the latent representation 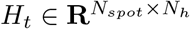.

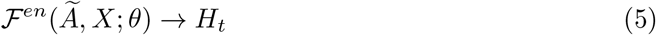

Subsequently, using the sampled subset ℰ_*drop*_, we construct the positive sample set 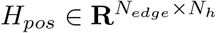, where *N*_*edge*_ represents the number of edges in the ℰ_*drop*_ subset. In detail, if the *i*-th spot and the *j*-th spot satisfy *e*_*ij*_ ∈ ℰ_*drop*_, then 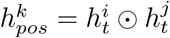. Where ⊙ represents the Hadamard product operation of vectors.

To generate the negative sample set, we randomly select spot pairs from all spots and create the 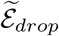 set. Each spot pair corresponds to a negative sample vector, denoted as 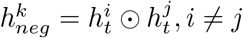. The probability of sampling negative spot pairs belonging to neighbors is very low due to the large number of spots compared to the number of nearest neighbors. We construct the negative sample set 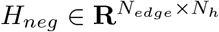. By training the relationship discriminator using positive and negative sample pairs, we improve its discriminability.

More precisely, the discriminator 𝒟 is composed of multiple layers of perceptrons and utilizes the Sigmoid function for final activation and normalization. Consequently, the output reflects the scores assigned to the positive and negative samples. The scoring computation is as follows:

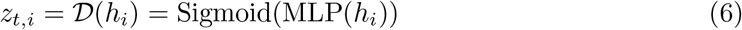

### Model training strategy

STMask combines gene representation learning and gene relationship learning through two key objectives. First, it enhances gene expression by reconstructing masked spot features. Second, it improves model discriminability by maximizing mutual information in the disrupted relationship network. To achieve these goals, STMask utilizes two loss functions, namely reconstruction loss and contrastive loss, for joint model optimization.

To reconstruct the masked features from the given partially observed input features, we utilize the Scaled Cosine Error (SCE) as an objective function. The *l*_2_-normalized cosine error enhances the stability of embedding representation learning. Under a predefined scaling factor *γ*, the similarity between the reconstructed prediction and the original input is calculated only on the masked nodes set. The math formulation is as follows:

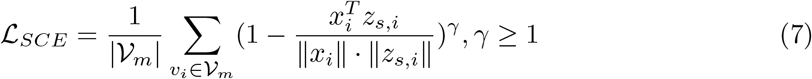

where *γ* is fixed at 2 throughout the entire experiment to reduce the weight of contributions from simple samples during training, and | 𝒱_*m*_| represents the number of nodes in the masked set.

Finally, Maximizing Mutual Information (MMI) serves as our objective for the contrastive loss:

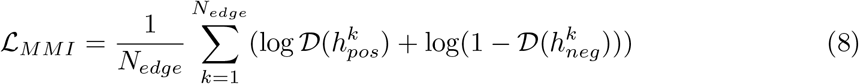

The entire learning objective is the weighted sum of the reconstruction loss ℒ_*SCE*_ and the contrastive loss ℒ_*MMI*_ :

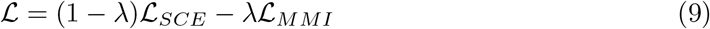

## Results

### Applying STMask to the DLPFC dataset

In this study, we selected five representative state-of-the-art methods, including two leveraging histological image features (SpaGCN [14], DeepST [16]), one based on maximizing mutual information (CCST [18]), and two deep autoencoder methods (SEDR [17], STAGATE [1]). We compared the Adjusted Rand index (ARI) and Normalized Mutual Information (NMI) of STMask with five currently representative methods (Table B in S1 Supplementary materials).

To quantitatively evaluate the spatial domain recognition performance of STMask, we first applied it to the DLPFC dataset. The results demonstrated that STMask excelled in clustering performance among the 12 tissue sections, exhibiting a remarkable median ARI value of 0.596. This represented a significant advancement of 6.9% and 6.5% compared to the respective median ARI values of 0.527 for STAGATE and 0.531 for SEDR. Furthermore, STMask also achieved the highest median NMI value on the DLPFC dataset (Fig 2B).

**Fig 1.**
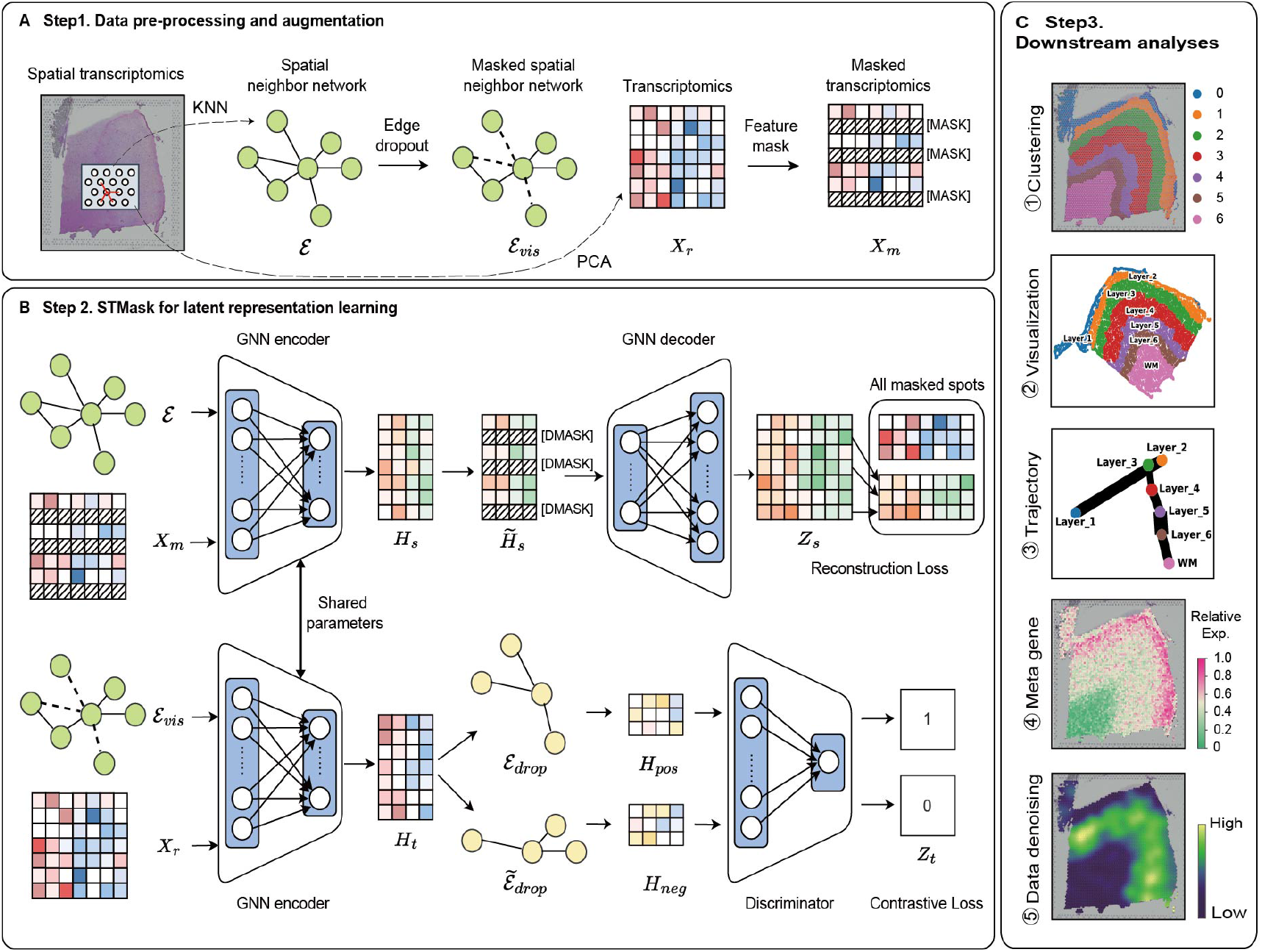
Overview of STMask. STMask employs two different masking techniques to handle the gene expression matrix and spatial topological structure, respectively. (B) The gene representation learning channel of STMask consists of GNN encoder-decoder pairs, using scaled cosine loss to reconstruct masked gene representations. The gene relationship learning channel of STMask comprises GNN encoder-discriminator pairs, leveraging the viewpoint of maximizing mutual information in contrastive learning to capture distinguishing features between positive and negative graph sample pairs. (C) The learned embeddings are utilized for spatial clustering, trajectory inference, gene expression imputation, and other downstream analysis tasks.

**Fig 2.**
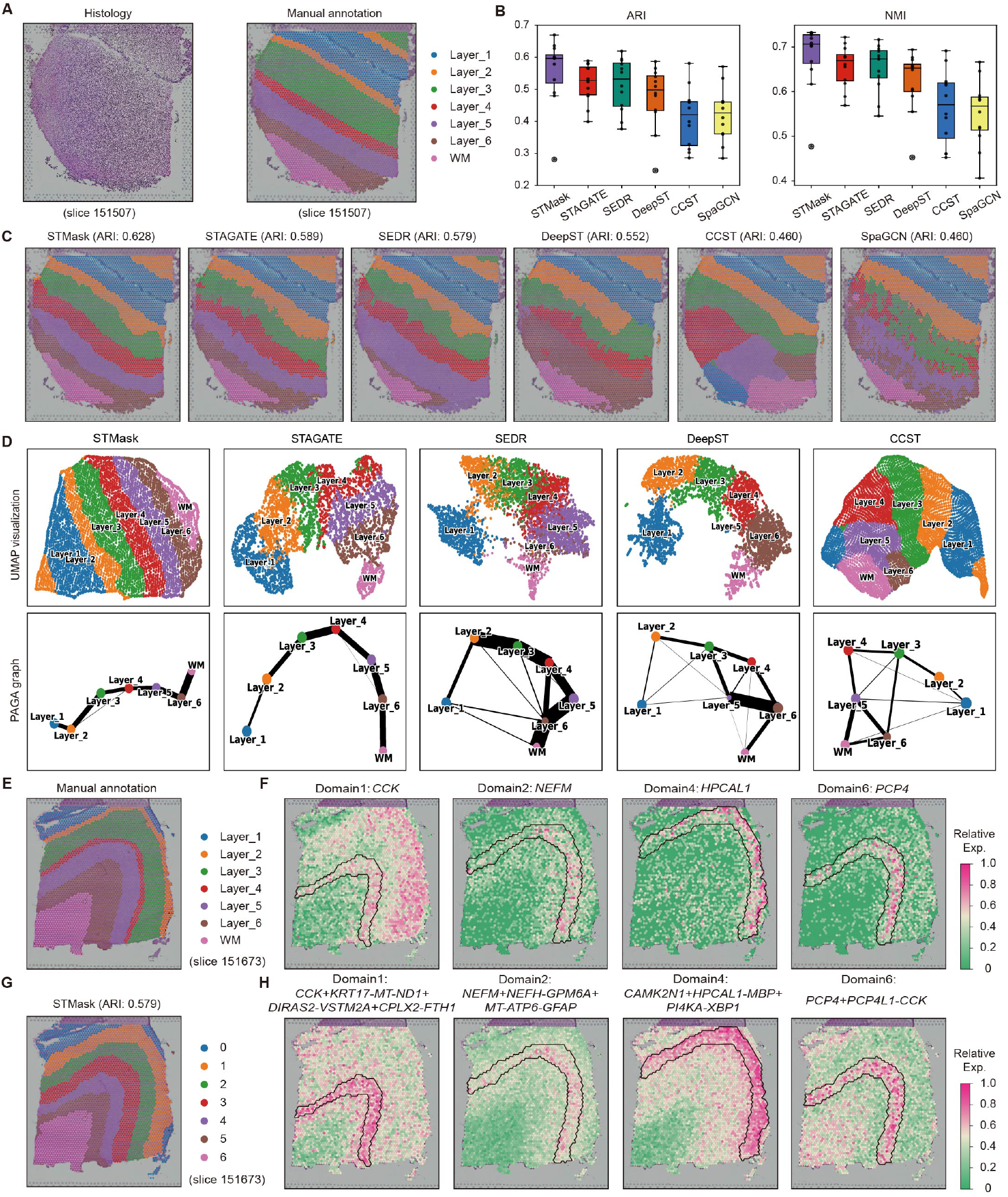
Identification of spatial domains and SVGs in the DLPFC dataset with STMask. (A) Tissue image (left) and manually annotated layer structures of the cortical layers and white matter (WM) on slice 151676 of the DLPFC (right). (B) Boxplots showing the clustering accuracy with ARI scores (left) and NMI scores (right) for six methods across all 12 slices of the DLPFC dataset. The center line, box, and whiskers represent the median, interquartile range, and 1.5 times the interquartile range, respectively. (C) Spatial domain identification on slice 151507 using STMask, STAGATE, SEDR, DeepST, CCST, and SpaGCN, respectively. (D) UMAP visualization and PAGA graph generated by embedding on slice 151507. (E) Manually annotated layer structures of slice 151673. (F) Spatial expression patterns of SVGs detected by STMask on slice 151673. (G) Domain identification using STMask. (H) Spatial expression patterns of meta-genes detected by STMask.

Taking the tissue slice 151507 from the DLPFC as an example (Fig 2A), containing 4,226 points and 33,538 genes [21], we compared the performance of STMask with existing methods in spatial domain recognition (Fig 2C). Observing that the STMask achieved the highest ARI score of 0.628, depicting clear layer boundaries and showing greater consistency with manually annotated spatial domains, thereby achieving optimal clustering accuracy. While SpaGCN utilized histological image features, it failed to distinguish between layers 4 and 6, presenting a significant mixture of spots across various domains, yielding unsatisfactory results. This could be due to its failure to extract features from histological images, and another reason could be that the model adopted a single unsupervised clustering approach, leaving considerable room for improvement and optimization. The results of CCST, utilizing mutual information maximization, demonstrated better performance in obtaining continuous domains, but failed to accurately identify the 4, 5, and 6 cortical layers, and the WM layer. This suggests the need for further exploration of gene features, as learning more expressive embeddings is crucial for domain recognition. Although DeepST effectively extracted histological information using deep neural networks, there were still mixtures of spots and discontinuous clusters in the spatial domain, with many outliers present. Analysis of the spatial domain recognition results revealed that STMask could effectively identify the expected cortical layer structures (Fig A in S1 Supplementary materials).

In the UMAP [26] plots generated from the embeddings by STMask (Fig 2D), the spatial relationships between each spatial domain were revealed, showing consistent spatial trajectories across layers (from layer 1 to 6 and white matter). This aligned with the functional similarity between adjacent cortical layers and followed a chronological order. Further confirmation of the inferred trajectories through PAGA [27] plots showed that only the embeddings from STMask and STAGATE exhibited the expected cortical layer structure and demonstrated nearly linear developmental trajectories. In contrast, the PAGA results from other baseline methods appeared mixed. Furthermore, we utilized an alignment objective function [28] to register multiple consecutive adjacent tissue slices through translation, rotation, and flipping for 3D tissue reconstruction, resulting in a significant improvement in the visual representation of 3D spatial patterns [2] (Figs B and C in S1 Supplementary materials).

We followed a similar procedure as in SpaGCN to detect spatially variable genes (SVGs) enriched in each spatial domain [14]. Differential expression (DE) analysis was conducted on spots within target and neighboring domains [29], selecting genes with an FDR-adjusted *P* value *<* 0.05 as SVGs. A total of 76 SVGs were detected on slice 151673. Among these, domain 0 accounted for 68 SVGs, while domains 1, 2, and 6 each had 1 SVG, and domain 4 had 5 SVGs. We represented the relative expression levels of relevant genes using different colors. For example (Fig 2F), *NEFM* was enriched in domain 2 (layer 3), *PCP4* in domain 6 (layer 5), and *HPCAL1* in domain 4 (layers 1 and 2), consistent with previously reported results [30]. When individual gene markers presented challenges in delineating expression patterns across certain neuronal layers, we constructed meta-genes composed of multiple genes to label the expression patterns of specific domains. For instance (Fig 2H), *CCK* exhibited relatively weak enrichment in layers 2, 3, and 6, with a relatively low number of spots. We explicitly enhanced the expression pattern on target domain 1 by integrating genes such as *KPT17, DIRAS2*, and *CPLX2* into a meta gene. Additionally, we successfully observed the transferability of these SVGs to several other tissue slices (Fig D in S1 Supplementary materials).

### Denoising gene expression with STMask on the DLPFC dataset

Raw ST data was prone to noise and dropout events caused by sequencing technologies. This hindered the accurate representation of underlying expression patterns, thus impacting downstream analyses such as cell clustering, differential expression analysis, and cell trajectory inference. To tackle this issue, we applied STMask to the DLPFC dataset. By constructing denoised and imputed gene expression matrices, we successfully reduced noise in the raw data and improved the identification of spatial expression patterns in genes.

On the 151674 slice of the DLPFC, we compared the expression profiles of six layer-marker genes (*VAT1L, PCP4, NEFH, CALB1, GNAL, CRYAB*) between the raw data and the denoised expression obtained using STMask (Fig 3A). While the raw gene expression exhibited a scattered pattern across layers for these six marker genes, the denoised gene expression distinctly revealed the layer-specific enrichments of these genes, a phenomenon validated against the Nissl data publicly available in the Allen Human Brain Atlas [31] (Fig 3B). Additionally, we observed an enrichment of the *PCP4* gene in layer 5, accompanied by a significant enhancement in its expression pattern, consistent with previous research findings [21, 30]. Further comparison of violin plots (Fig 3C and 3D) between the raw and imputed data revealed an enhancement in the spatial expression patterns of the layer-marker genes, aligning more closely with manually annotated tissue structures (Fig E in S1 Supplementary materials). The experimental findings underscore the effectiveness of STMask in reducing noise while preserving spatial expression patterns.

**Fig 3.**
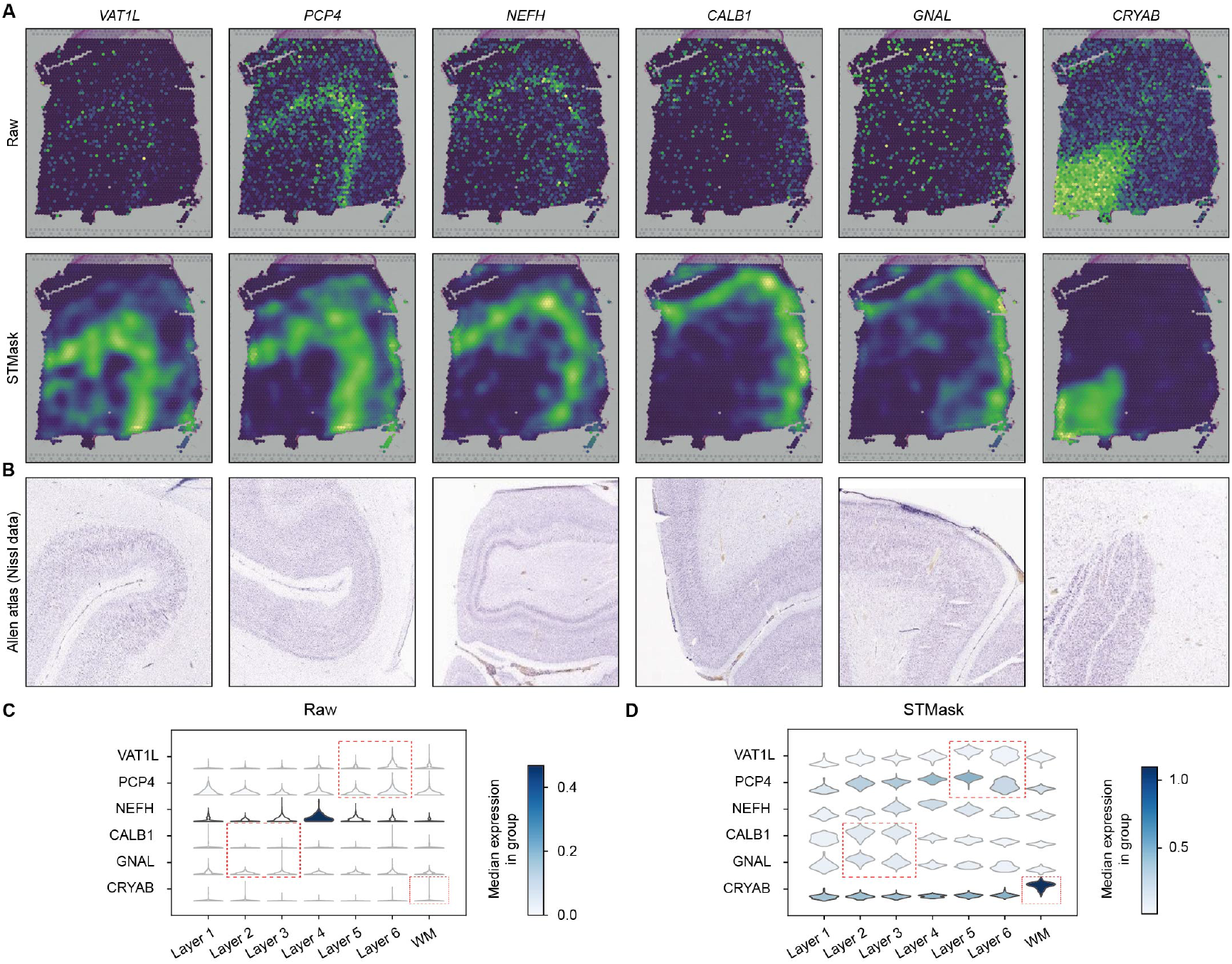
Enhancement of spatial patterns and resolution of layer-specific marker genes in the DLPFC Dataset by STMask. (A) Visualization of the original spatial expression and denoised data by STMask for six layer-specific marker genes on slice 151674 of the DLPFC. (B) Nissl images from the publicly available Allen Human Brain Atlas. (C) Violin plots of the original expression of layer-specific marker genes. (D) Violin plots of the denoised expression of layer-specific marker genes by STMask. The cortical layers corresponding to the layer-specific marker genes are highlighted with a red box.

### Applying STMask to the BRCA dataset

We analyzed the BRCA dataset with 20 regions, including DCIS/LCIS, healthy tissue, IDC, and low malignant tumor margins (Fig 4A). Due to the significant differences between the main tissue regions in BRCA histological images, DeepST and SpaGCN, leveraging histological information, achieved decent performance (Fig 4B). However, the identified spatial domains still exhibited blurry boundaries and numerous outliers (Fig 4C). STAGATE and SEDR failed to accurately identify the invasive ductal carcinoma regions (IDC 2/4/5) and erroneously fragmented them into many small clusters. Compared to existing methods, STMask, although independent of histological images, still attained the highest clustering performance scores. It demonstrated clear and smooth tissue boundaries and fewer outliers in spatial domain identification.

**Fig 4.**
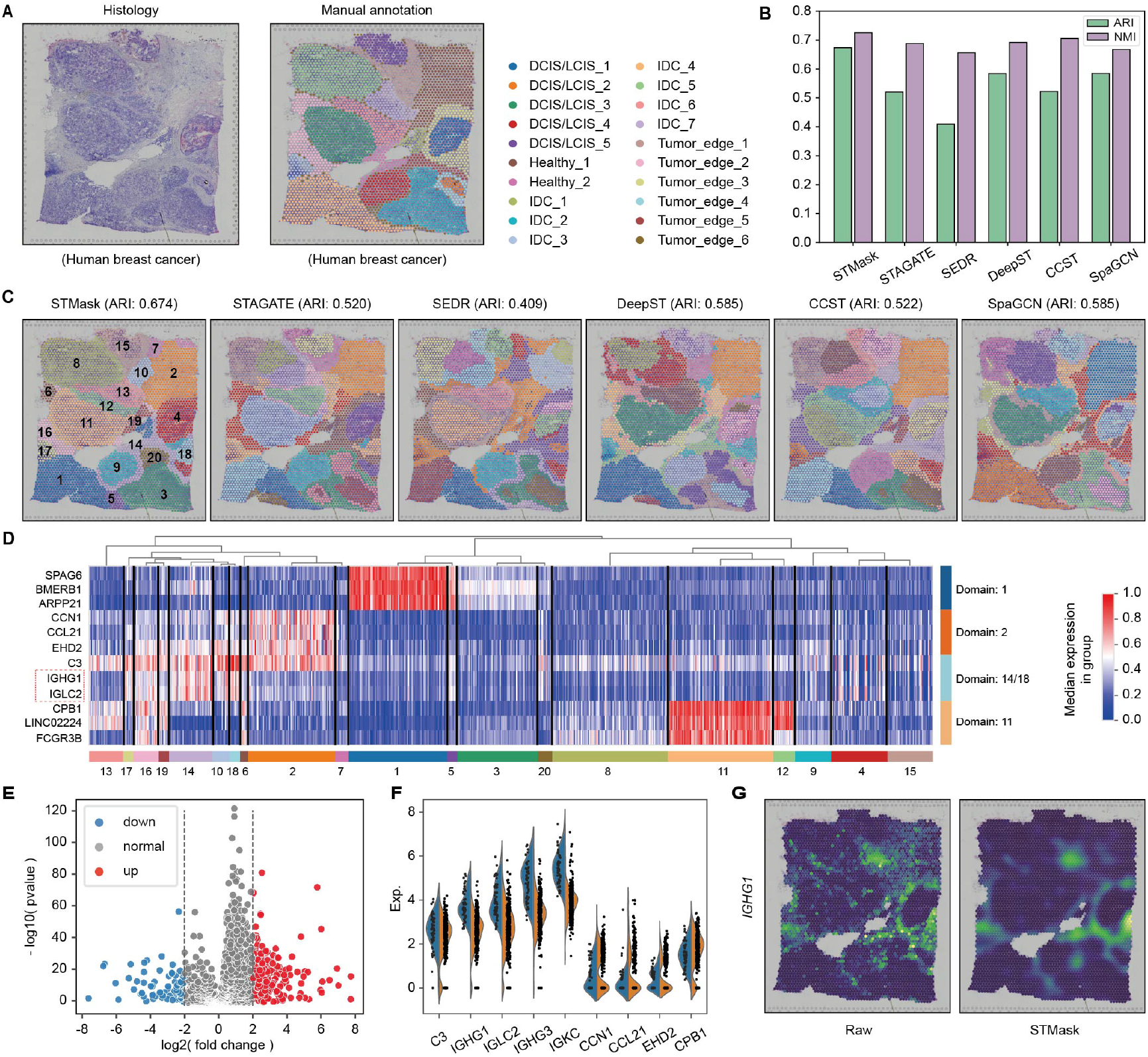
Experiment results on the BRCA dataset. (A) Tissue images of BRCA (left) and manually annotated layer structures (right). (B) Par chart illustrating clustering accuracy across various methods on BRCA, measured by ARI and NMI scores. (C) Spatial domain recognition for BRCA. (D) Heatmap illustrating the expression of structural domains on the top three DEGs from clusters 1, 2, 14/18, and 11. (E) Differential expression analysis between clusters 2 and 18. (F) Volcano plot of DEGs between clusters 2 and 18. (G) Visualization of the original spatial expression and expression after denoising with STMask using *IGHG1* as layer marker genes.

To dissect the heterogeneity of tumor regions, we compared the expression of the top three Differentially Expressed Genes (DEGs) in cluster 1 (IDC), cluster 2 (healthy), cluster 14/18 (tumor edge), and cluster 11 (DCIS/LCIS) across all domains and visualized their expression patterns with a heatmap to understand the relationship between gene expression and clusters (Fig 4D). The results revealed significant heterogeneity among these clusters. The genes *IGHG1* and *IGLC2* were predominantly enriched in cluster 14/18, while *CCN1* and *CCL21* were mainly enriched in cluster 2. Additionally, we conducted differential expression analysis between cluster 18 and cluster 2 to further explore the differences in gene expression between tumor and healthy tissues, detecting a total of 282 DEGs (|*log*2*FoldChange*| ≥ 2 and *Pvalue <* 0.05) (Fig 4E). Violin plots displayed the distribution of nine high-ranking DEGs in the two clusters (Fig 4F). Among these crucial DEGs, the expression of the immunoglobulin heavy constant gamma 1 (*IGHG1*) gene was positively correlated with immunoglobulin levels originating from various tumor sources, participating in pathological processes such as tumor cell proliferation, migration, invasion, immune evasion, and epithelial-mesenchymal transition. Research suggested that *IGHG1* promotes malignant development of breast cancer by activating the AKT pathway [32] and may serve as an effective target for breast cancer treatment. Finally, we demonstrated the denoising performance of STMask (Fig F in S1 Supplementary materials). The denoised gene expression by STMask better reflects the enrichment of layer-marked genes and spatial expression patterns compared to the raw. For example, using *IGHG1* as a layer marked gene (Fig 4G), after denoising, there was a noticeable enrichment of *IGHG1* in clusters 10, 14, 16, and 18 associated with tumor edges, which is more consistent with manual annotation of tissue structure.

### Applying STMask to the HM dataset

We analyzed the HM dataset (Fig 5A), which includes melanoma, stroma, lymphoid tissue, and an additional unannotated region. Bar plots (Fig 5B) showed that STMask had the best clustering performance, while CCST had the lowest due to over-smoothing with its four-layer GCN backbone on the sparse and low-density HM dataset. Spatial domain recognition plots (Fig 5C) revealed that STMask accurately identified melanoma, stroma, and lymphoid tissue regions with clear boundaries, while other methods failed to identify stroma.

**Fig 5.**
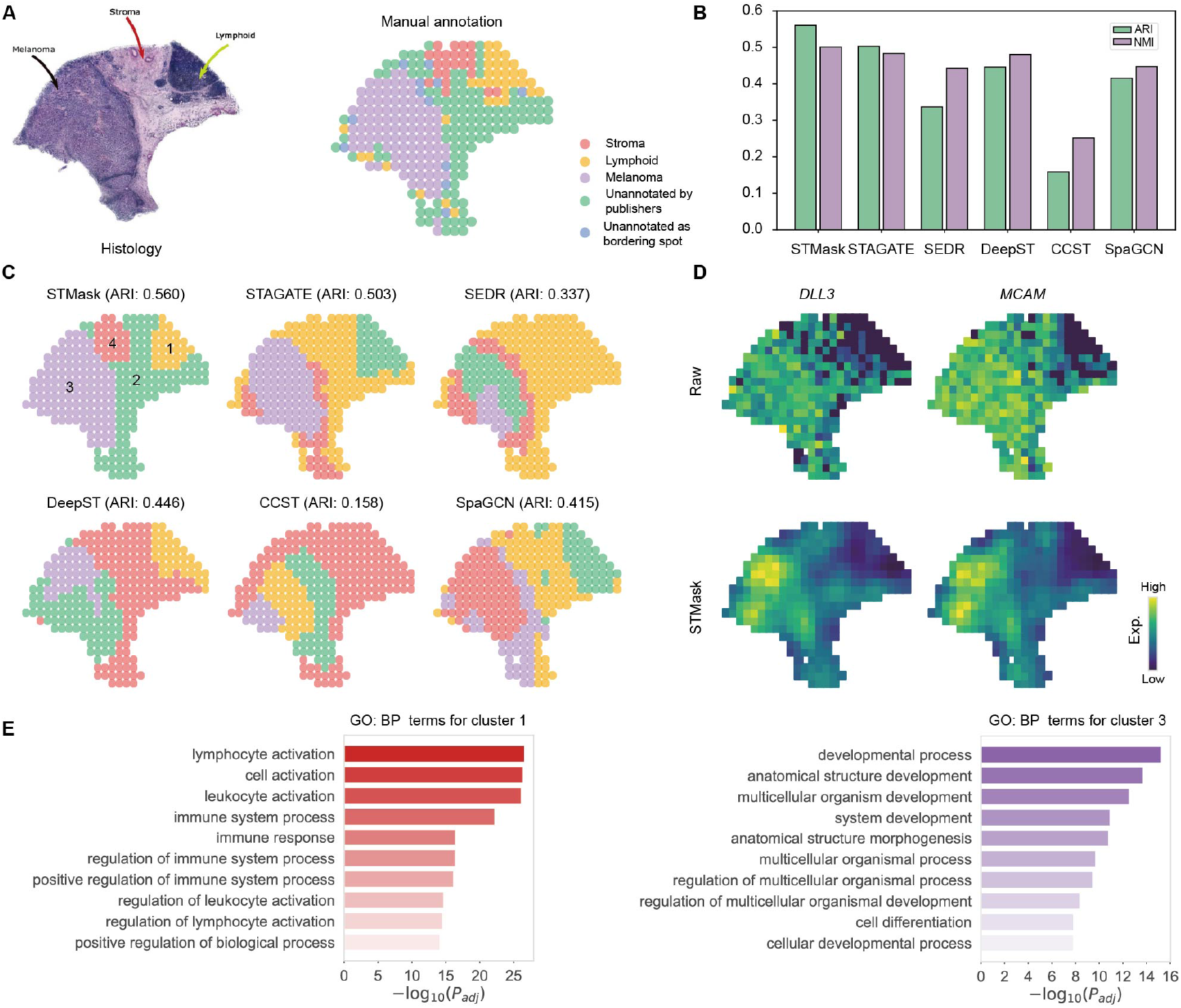
Performing spatial domain recognition in the HM dataset. (A) Tissue image of melanoma (left) and manually annotated layer structures (right). (B) Bar chart illustrating clustering accuracy across various methods on the melanoma dataset, measured by ARI and NMI scores. (C) Spatial domain recognition for HM. (D) Visualizing the original spatial expression and the expression after denoising with STMask using *DLL3* and *MCAM* as layer marker genes. (E) Here are the top 10 significant GO:BP terms for Cluster 1 (Lymphoid) and Cluster 3 (Melanoma) after imputation.

Continuing, we conducted differential expression analysis for cluster 1 and cluster 3 to further investigate the differences in gene expression (|*log*_2_(*FoldChange*)| ≥ 2 and *P value <* 0.05). These DEGs played crucial roles in various physiological and pathological processes. For instance, delta-like ligand 3 (*DLL3*) was highly expressed in some neuroendocrine tumors while being rarely expressed in normal tissues, providing potential for targeted therapy [33]. Additionally, related studies indicated that the melanoma cell adhesion molecule (*MCAM*) was involved in the adhesion and metastasis processes of melanoma cells, exhibiting different expression patterns in various types of melanoma and showing a close correlation with the malignancy and metastatic capacity of melanoma [34]. After denoising the genes, we observed an enrichment of *DLL3* and *MCAM* genes in the melanoma domain, consistent with previous research findings (Fig 5D and Fig G in S1 Supplementary materials). Finally, we conducted functional enrichment analysis for cluster 1 and cluster 3 using the Gene Ontology: Biological Process (GO:BP) database [35]. Under the same conditions, gene expression imputed by STMask obtained more GO:BP terms. The identified cluster 1 corresponded to the lymphoid family, with the displayed 10 GO:BP terms mostly associated with changes in cell morphology and behavior, as well as the maintenance or suppression of immune responses. Cluster 3 corresponded to the melanoma family, and its abundant terms were mostly related to biological regulation and metabolic processes (Fig 5E).

### Applying STMask to the MBA dataset

Finally, we analyzed the MBA dataset. Compared to the DLPFC dataset, it exhibits a more intricate tissue structure, posing greater challenges. We considered 52 clusters as recognizable spatial domains and compared STMask with existing methods in spatial domain recognition based on clustering performance. From the bar graph displaying ARI and NMI scores, it was evident that STMask achieved the highest scores. STAGATE and SpaGCN mistakenly partitioned the central nucleus (CPu) into multiple regions, while DeepST and SpaGCN exhibited noticeable spot mixing, resulting in unclear boundaries between clusters (Fig H in S1 Supplementary materials).

### Ablation studies

To further investigate the contributions of each component of STMask to spatial domain recognition, we conducted a series of ablation experiments on the DLPFC, BRCA, HM, and MBA datasets. Initially, we explored the impact of the masking mechanism on clustering performance in each channel of the dual-channel autoencoder. We designed three variants of STMask. The STMask **w/o Delete Edge** variant only masked gene expression in the gene representation learning channel, removing the masking operation on adjacency relationships in the gene relationship learning channel. Conversely, the STMask **w/o Mask Node** variant retained masking on spot-to-spot relationships in the gene relationship channel while removing masking operations on gene expression for spots. The STMask **Neither** variant was an overcomplete autoencoder that did not utilize any masking mechanisms to prevent overfitting. Table 2 presented the average ARI and NMI clustering scores of STMask and its variants over 10 runs on the four datasets and Fig 7A displayed a bar chart showing the ARI results of STMask and its variants across 12 slices of DLPFC. STMask outperformed all variants in terms of ARI, except for slightly lower NMI scores compared to the w/o Delete Edge and w/o Mask Node variants on the MBA dataset. Notably, STMask Neither, which did not utilize masking mechanisms, performed the worst across all datasets. Thus, this experiment strongly demonstrated the competitiveness of graph masking autoencoders in spatial domain clustering, with the dual-channel graph masking autoencoder showing significant performance improvement, demonstrating true effectiveness in spatial domain recognition.

**Table 2.** Ablation studies on the contribution of masking mechanism and GNN type to model clustering performance. Considering that the DLPFC dataset comprises 12 tissue slices, we used the median clustering metrics (ARI, NMI) of these slices to represent the overall performance.

| Dataset | DLPFC |  | BRCA |  | HM |  | MBA |  |
| --- | --- | --- | --- | --- | --- | --- | --- | --- |
|  | ARI(%) | NMI(%) | ARI(%) | NMI(%) | ARI(%) | NMI(%) | ARI(%) | NMI(%) |
| STMask | **59.39±0.12** | **70.59±0.11** | **69.06±1.48** | **72.92±0.31** | **57.41±9.37** | **53.82±5.40** | **46.22±4.20** | 70.36±0.52 |
| w/o Delete Edge | 55.69±1.36 | 69.19±1.11 | 62.30±1.79 | 71.95±0.71 | 38.77±3.64 | 38.34±2.61 | 36.12±1.32 | **73.20±0.54** |
| w/o Mask Node | 56.25±0.79 | 69.21±0.63 | 59.91±2.86 | 71.46±0.82 | 31.72±2.23 | 38.67±0.58 | 38.39±0.66 | 72.51±0.21 |
| Neither | 45.53±0.11 | 61.85±0.06 | 44.32±1.47 | 63.03±0.24 | 27.78±5.53 | 37.81±3.67 | 39.38±3.25 | 70.34±0.63 |
| GCN | **59.39±0.12** | **70.59±0.11** | **69.06±1.48** | **72.92±0.31** | **57.41±9.37** | **53.82±5.40** | **46.22±4.20** | 70.36±0.52 |
| GAT | 54.09±1.64 | 68.03±0.36 | 60.81±1.99 | 71.09±0.81 | 37.97±6.04 | 38.77±3.16 | 38.41±1.12 | 73.01±0.26 |
| FeaSt | 53.31±0.39 | 67.52±0.31 | 56.83±2.76 | 70.66±0.79 | 48.51±0.65 | 48.58±0.70 | 38.82±1.31 | 73.40±0.59 |
| Transformer | 54.77±0.33 | 68.93±0.55 | 62.47±1.08 | 71.70±1.00 | 45.94±2.85 | 45.85±1.91 | 37.88±0.73 | **73.55±0.24** |
| ResGatedGraph | 52.59±0.98 | 67.45±0.25 | 61.64±2.50 | 70.91±1.11 | 43.02±3.32 | 43.59±1.27 | 35.83±2.30 | 71.78±0.84 |
| SAGE | 54.07±1.15 | 68.94±1.00 | 58.85±1.86 | 70.80±0.84 | 45.48±5.28 | 45.95±3.03 | 38.22±1.02 | 73.19±0.38 |

Furthermore, we investigated the impact of different masking ratios in the dual channel on model performance. By employing a grid search approach, we determined the optimal masking rate parameters (feature mask ratio *ρ*_*m*_ and edge mask ratio *ρ*_*d*_). We visualized the ARI scores of STMask on the four datasets with heatmaps (Fig 6), where higher brightness indicated higher ARI values, and conversely, lower brightness indicated lower ARI values. On the BRCA and HM datasets, a pattern emerged with higher brightness in the lower-left region and lower brightness in the upper-right region (Fig 6B and 6C), suggesting a similar contribution of dual-channel masking to model performance. A general pattern of higher brightness on the left and lower brightness on the right was observed on the DLPFC and MBA datasets (Fig 6A and 6D), indicating a significant impact of masking rates in the gene relationship learning channel on model performance, while being less sensitive to the magnitude of feature masking rates. Combining the results of the ablation experiments on masking mechanisms and these observations, we can conclude that the dual-channel masking self-supervised learning method has made significant contributions to spatial domain recognition in ST data.

**Fig 6.**
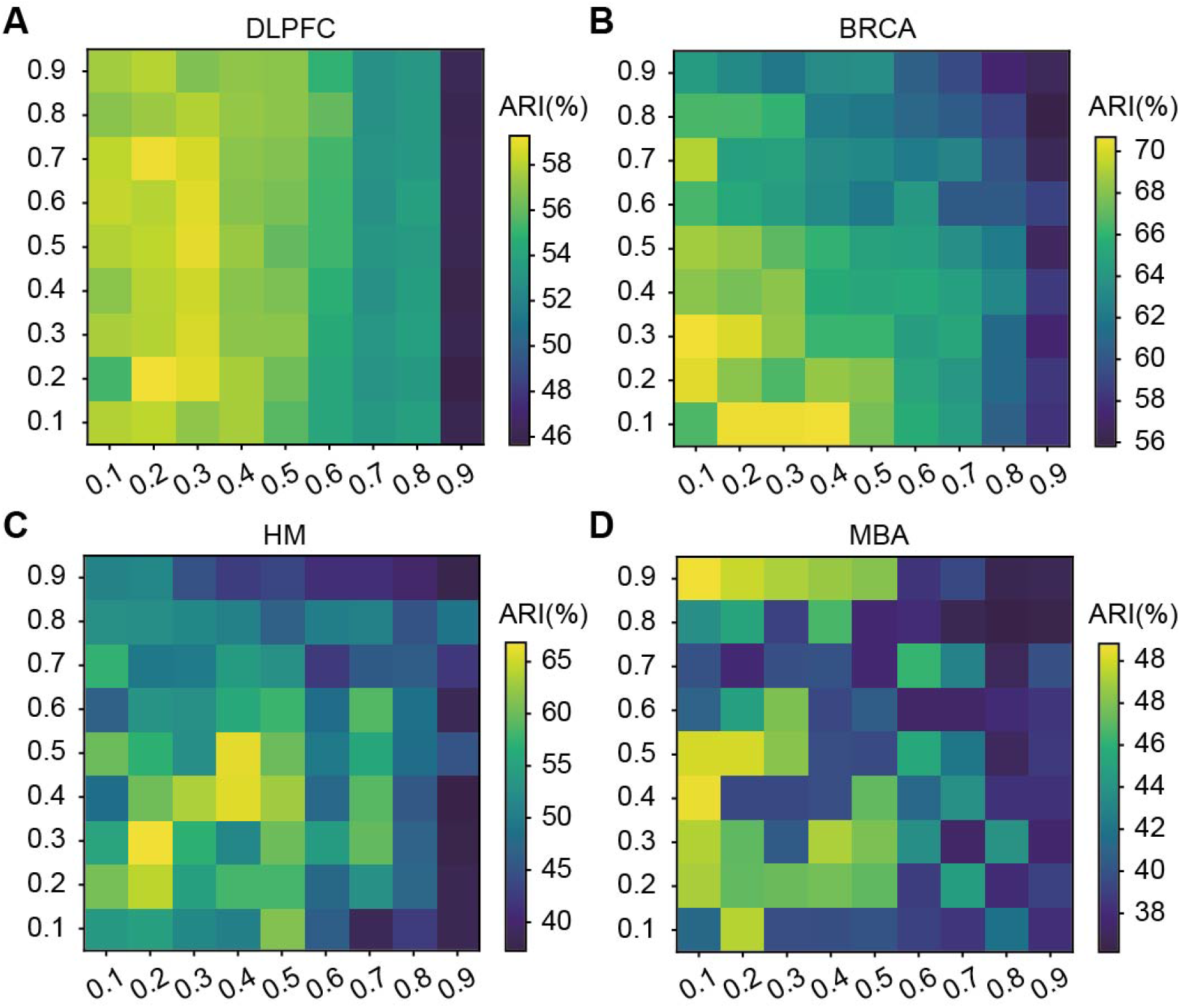
Ablation studies of mask ratio for STMask on DLPFC, BRCA, HM and MBA datasets. In each heatmap, the horizontal axis values represent the edge mask ratio (*ρ*_*d*_), and the vertical axis values represent the feature mask ratio (*ρ*_*m*_).

**Fig 7.**
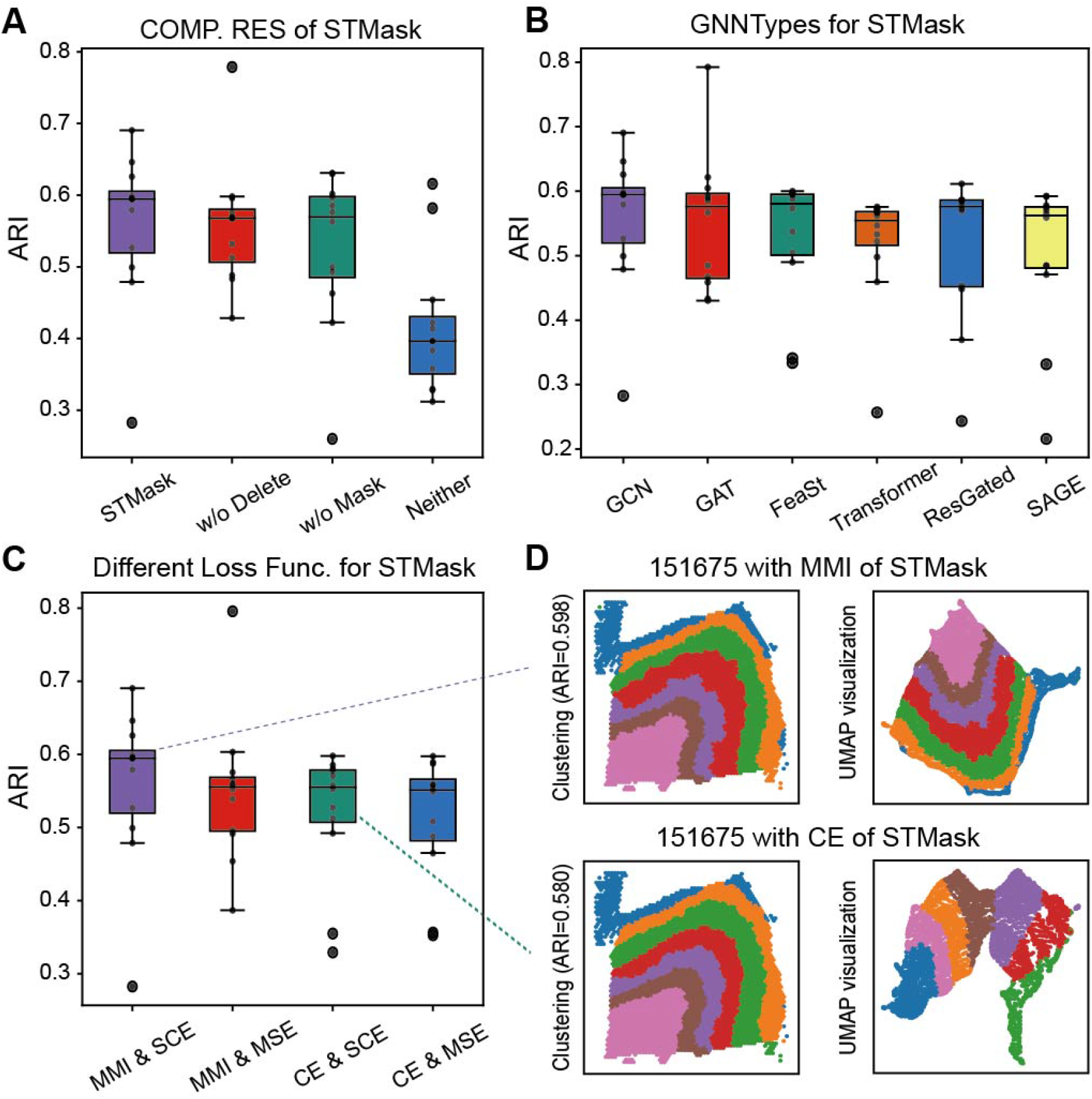
Ablation studies on the components. Including GNN types, and objective functions on the DLPFC dataset.

We also explored the impact of different GNN Types (including GCNConv, GATConv, FeaStConv, TransformerConv, etc.) as the backbone of graph autoencoders on the overall model performance. Table 2 presents the average ARI and NMI evaluation scores obtained from running these models 10 times across four datasets. From the results, it was evident that when GCN serves as the backbone of the autoencoder, the model performs exceptionally well on each dataset and exhibits good robustness. Furthermore, we ran six GNN types on 12 DLPFC slices and computed their respective ARI values (Fig 7B). In the best-case scenario, they all demonstrated decent performance, with GCNConv and GATConv achieving relatively higher ARI values. This experiment underscores that the model performance is minimally affected by the GNN type, while the dual-channel masking mechanism remains a key factor in significantly enhancing clustering performance. Finally, we investigated the impact of different objective functions on model performance on the DLPFC dataset, demonstrating that the combination of SCE and MMI yielded promising results (Fig 7C).

## Discussion

In this paper, we propose an efficient self-supervised learning method STMask utilizing a dual-channel masked graph autoencoder for the analysis and research of ST data. The gene representation learning channel cleverly integrates the masking mechanism into the graph autoencoder, reducing feature redundancy and enhancing model robustness. Additionally, we employ the scaled cosine loss as the training objective to stabilize the embedding representation. Simultaneously, the gene relationship learning channel disrupts the spatial graph structure to generate multiple perspectives and combines contrastive learning principles to maximize mutual information, thereby strengthening the model’s discriminative capability. It is worth noting that the two channels share the same encoder, which helps to enhance the model’s ability to learn gene expressions and facilitates the acquisition of discriminative feature embeddings.

In spatial domain recognition, STMask shows clearer boundaries, fewer mixed spots, and spatial domains more consistent with manual annotations, further detecting spatially variable genes with rich expression patterns within the identified domains. This is mainly attributed to STMask cleverly integrating a masking mechanism into a self-supervised graph model, compelling the masked gene expressions to learn more robust representations from neighbors. Additionally, STMask explores the correlation between gene expression and spatial information by maximizing mutual information. Through denoising experiments, we further explore the importance of denoised gene expression in identifying biologically relevant domains. In breast cancer data, we detect various differentially expressed genes (e.g., *IGHG1*) predominantly enriched in tumor regions, which may serve as effective targets for cancer therapy, providing crucial insights into its pathobiological processes. Lastly, ablation studies highlight the significant contributions of the dual-channel and masking mechanisms in STMask, as well as the impact of different masking ratios on clustering performance. By adeptly leveraging masking mechanisms and integrating contrastive learning approaches, STMask can effectively identify spatial domains and integrate spatial and feature information.

While STMask performs well in spatial domain recognition, it has yet to effectively address the batch effects across multiple tissue slices. We plan to introduce discriminative loss among multiple tissue slices to make the masking mechanism more versatile and effective in handling and analyzing ST data.

## Supporting information

**S1 Supplementary materials. Supplementary figures, tables and analysis details**. Details on comparison with other spatial domain identification methods, STMask parameter settings, evaluation citeria, detection of SVGs and spatially variable mete genes and alignment of consecutive ST slices. **Fig A. Spatial domain recognition on 12 DLPFC slice datasets**. Comparison of spatial domains by clustering assignments via STMask, STAGATE, SEDR, DeepST, CCST, SpaGCN, and manual annotation in all 12 sections of the DLPFC dataset. **Fig B. UMAP visualization and PAGA graphs on 12 DLPFC slice datasets**. UMAP visualization and PAGA graphs generated by STMask, STAGATE, SEDR, DeepST and CCST embeddings respectively. Spots are colored by their manual annotations. **Fig C. Consecutive slices of a tissue for 3D slice alignment**. (A) The spatial domain recognition results for consecutive slices 151507-151510, aligned in 2D (middle), and displayed as a stack of the four slices in 3D (right). (B) Consecutive slices 151669-151672. (C) Consecutive slices 151673-151676. **Fig D. Detection of SVGs and spatially variable mete genes**. (A) The five spatially variable genes detected by STMask on slices 151507 and 151673. (B) The spatially variable meta genes detected by STMask on slices 151507 and 151673. **Fig E. Comparison of the spatial expression patterns before and after STMask denoising**. (A) Ground-truth segmentation of cortical layers and white matter (WM) in the DLPFC section 151507. (B) Expression visualization of five layer-marker genes in the DLPFC section 151507. (C) and (D) Violin plots of the raw expressions (C) and the STMask denoised expressions (D) of layer-marker genes in the DLPFC section 151507 respectively. The cortical layer corresponding to the layer-marker genes is marked with red boxes. **Fig F. Experiment results in the BRCA dataset**. (A) Manually annotated BRCA. (B) Volcano plot of DEGs between DCIS/LCIS region and IDC region. (C) Differential expression analysis between DCIS/LCIS region and IDC region. (D) UMAP visualization generated through embedding. (E) Expression visualization of five layer-marker genes in the BRCA dataset. **Fig G. Experiment results in the HM dataset**. (A) Tissue images of HM. (B) UMAP visualization generated through embedding. (C) Expression visualization of five layer-marker genes in the HM dataset. (D) Violin plots of the STMask denoised expressions of layer-marker genes. (E) Volcano plot of DEGs between lymphoid region and melanoma region. (F) Differential expression analysis between lymphoid region and melanoma region. **Fig H. Experiment results in the MBA dataset**. (A) Manually annotated MBA. (B) Bar chart illustrating clustering accuracy across various methods on the MBA, measured by ARI and NMI scores. (C) Spatial domain recognition for MBA. (D) UMAP visualization generated through embedding. **Table A. Summary of the datasets in this study. Table B. Summary of the 5 clustering methods based on methodology, algorithm input and code link**.

## Data availability

The datasets used in this paper can be downloaded from the following websites. Specifically, (1) The LIBD human dorsolateral prefrontal cortex (DLPFC) dataset (http://spatial.libd.org/spatialLIBD/); (2) 10x Visium ST dataset of human breast cancer (BRCA) (https://support.10xgenomics.com/spatial-gene-expression/datasets/1.1.0/V1_Breast_Cancer_Block_A_Section_1); (3) The human melanoma (HM) dataset obtained through the ST platform (https://www.spatialresearch.org/resources-published-datasets/doi-10-1158-0008-5472-can-18-0747/); and (4) 10x Visium spatial transcriptomics dataset of mouse brain sagittal-anterior (MBA) (https://www.10xgenomics.com/datasets/mouse-brain-serial-section-1-sagittal-anterior-1-standard-1-0-0) and the Allen Mouse Brain Reference Atlas (https://mouse.brain-map.org/static/atlas). The above datasets are available on https://github.com/donghaifang/STMask.

## Author contributions

**Conceptualization**: Wenwen Min, Donghai Fang, Jinyu Chen, Shihua Zhang.

**Funding acquisition**: Wenwen Min and Jinyu Chen.

**Investigation**: Wenwen Min, Donghai Fang, Jinyu Chen, Shihua Zhang.

**Methodology**: Wenwen Min, Donghai Fang.

**Project administration**: Wenwen Min.

**Resources**: Wenwen Min, Donghai Fang.

**Software**: Wenwen Min, Donghai Fang.

**Supervision**: Wenwen Min.

**Validation**: Wenwen Min, Donghai Fang.

**Visualization**: Wenwen Min, Donghai Fang.

**Writing–original draft**: Wenwen Min, Donghai Fang.

**Writing–review & editing**: Wenwen Min, Donghai Fang, Jinyu Chen, Shihua Zhang.

